# Distribution patterns of two widespread, co-dominant savanna tree species of the Indian subcontinent under current and future climates

**DOI:** 10.1101/2025.07.13.664606

**Authors:** Kaikho Liriina, Biswa Bhusana Mahapatra, Abhishek Gopal, Aravind Madhyastha, Jayashree Ratnam, Mahesh Sankaran

## Abstract

Future climate is expected to alter the dynamics and distribution of vegetation. Here, we use ecological niche modelling to investigate the current and predicted future distributions of two dominant tree species that characterize the ecology and dynamics of the widespread tropical dry biomes of the Indian subcontinent- *Terminalia anogeissiana* (formerly and henceforth *Anogeissus latifolia)* and *Terminalia elliptica* (both Combretaceae). Using current occurrence data and six ecologically relevant climate variables, we find that, as expected, *T. elliptica* and *A. latifolia* share highly similar climatic niches with substantial overlap (∼84%). Mean annual precipitation (MAP), mean temperature of the driest quarter (MTDQ), and temperature annual range (TAR) emerge as the key environmental variables explaining the distribution of both species, but their relative influence differs. *T. elliptica* appears to occur across a broader climatic niche that includes wetter areas with cooler dry seasons (lower MTDQ) and lower temperature annual ranges (TAR) relative to *A. latifolia.* Currently, the area of modelled suitable habitat for both species are vast (*A. latifolia*: 416,057 km²; *T. elliptica*: 401,544 km²). However, under future climate scenarios, both species are projected to incur dramatic losses of suitable habitat (*A. latifolia*: ∼63% by 2041-2060 and ∼84% by 2081-2100; *T. elliptica*: ∼57.8% by 2041-2060 and ∼73% by 2081-2100). By 2100, the Western Ghats are projected to remain as the last key refugia for both species, while vast other regions are likely to become increasingly unsuitable. Contrary to current understanding, our results underscore the high vulnerability of currently dominant tree species of the tropical dry biomes, through potential losses of suitable habitats under future climate regimes.

## 1. INTRODUCTION

Climate is a major factor determining the distribution of biomes worldwide, with specific climatic regimes being associated with particular plant communities or functional types [1–3]. However, with the Earth’s climate changing rapidly, anticipated shifts - such as rising temperatures, increased rainfall, altered soil moisture, and prolonged dry seasons - are likely to have significant impacts on vegetation communities [4]. Global climatic changes are predicted to alter species distributions, interactions and associations, as well as community composition, potentially leading to species extinctions and large-scale changes in the patterns and distribution of global biodiversity [5]. Tropical dry biomes, including thorn and scrub forests, savannas and deciduous forests, are projected to be especially sensitive and vulnerable to these future climatic changes [6–11]. A detailed understanding of how plants in these important biomes will respond to these future changes is, thus, a prerequisite for effective conservation and mitigation efforts.

Tropical dry biomes are amongst the most widespread ecosystem types globally, and also in the Indian subcontinent [12–16]. They support a diverse and unique array of species, including many plant groups that are confined to this biome [14,17,18], distinct and unique faunal assemblages, as well as the nature-based livelihoods of millions of humans who live in these regions [14,16]. They are also amongst the most threatened ecosystems globally, with nearly 97% of their remaining area estimated to be at risk from a range of factors including land use and climate change [9,12]. However, unlike their rainforest counterparts, tropical dry forests have received much less attention in the literature, particularly in the Asian context, and our understanding of how these biomes are likely to respond to future changes in climate remains inadequate [9,14].

In this study, we examine the current and potential future distributions of two co-dominant and widespread tree species characteristic of tropical broad-leaved dry deciduous forests and mesic savannas of India -*Terminalia anogeissiana* (formerly *Anogeissus latifolia;* retained as such and henceforth *A. latifolia* in this manuscript) and *Terminalia elliptica* - both of which belong to the Combretaceae family [19–21]. We chose to focus on these two co-dominant species as dominant species are well recognized to play crucial roles in maintaining ecosystem functioning, stability and resilience [22,23]. For instance, both *A. latifolia* and *T. elliptica* have been reported to significantly influence primary productivity [24,25] and nutrient cycling [26,27], and contribute to carbon sequestration and storage in the dry biomes of the Indian sub-continent [28–31]. In addition, they also play key roles in regulating species diversity and composition, and resistance and resilience to disturbances such as fire [32]. Despite their ecological significance, both *A. latifolia* and *T. elliptica* are currently threatened by overexploitation for fodder, fuel, timber, medicine, and tannin extraction [33], along with chronic anthropogenic disturbances that hinder their recovery [34]. These species are also sensitive to extreme environmental stressors, with historical records showing large-scale mortality during prolonged droughts [35,36] and fire [37], highlighting their vulnerability to environmental changes. Given that shifts in the dominance of these species can substantially influence ecosystem dynamics, understanding their potential future distributions can provide valuable insights into ecosystem functioning and serve as early warning signs of fluctuations in ecosystem stability due to environmental changes.

Here, we used niche-modelling to project the current and future distributions of both species under different climate change scenarios. Niche-based models, such as species distribution models (SDMs) or environmental niche models (ENM), are quantitative tools that combine species occurrence data with environmental variables to identify suitable habitats and project future distributions under different climate scenarios [38–40]. These models are extensively used to quantify climatic niches of species [41,42], and to assess the impacts of global climate change on species distributions [43–45].

Our primary objectives were: a) to model the current distribution patterns of *A. latifolia* and *T. elliptica* based on key climatic predictors, and identify the key environmental factors influencing their distribution and occurrence patterns; b) to assess the niche spaces of *A. latifolia* and *T. elliptica*, anticipating that while their ranges may overlap, each species will have distinct habitat preferences; and c) to project their future distributions under different climate change scenarios, with expectations of increased temperatures and erratic rainfall patterns potentially leading to contractions of suitable habitat for both species.

## 2. METHODS

### 2.1 Study species

*Anogeissus latifolia* and *Terminalia elliptica* are closely-related species within the Combretaceae family. *Anogeissus latifolia* has been renamed *Terminalia anogeissiana* in recognition of its phylogenetic position within the *Terminalia* genus [46]. However, we retain the earlier nomenclature in this manuscript to enable its linkage to a large body of earlier literature that discusses the ecology and distribution of the species, which is highly relevant to the questions posed in this study.

*Terminalia elliptica* is a complex that includes *T. crenulata*, *T. coriacea*, *T. alata* var. *alata*, and *T. alata* var. *nepalensis* [47], all of which were subsumed under a single species *Terminalia tomentosa* and subsequently revised as *Terminalia elliptica* (Plants of the World) [48]. The *T. elliptica* complex is distributed widely across South and Southeast Asia, whereas *A. latifolia* is a single species, and is restricted to the Indian subcontinent. Both species range across the sub-Himalayan regions of the Indian subcontinent, including the Siwalik in the north and Aravali ranges in the west of the country, the Vindhya and Satpura ranges of the central Indian highlands, and southward throughout much of the Indian Peninsula [35,49]. They occur as co-dominants across large parts of their range, forming repeatable *Anogeissus-Terminalia* associations [19,34,50–52]. However, they also occur as monodominant stands in certain regions [34]. Both species are deciduous in nature, drought-tolerant and require abundant light, and are typically associated with dry environments, with a few species extending into moderately moist areas [35].

### 2.2 Occurrence data

Species occurrence data were collated from multiple open-source platforms, including the Global Biodiversity Information Facility (GBIF) [53], the India Biodiversity Portal (IBP) [54], citizen science-season watch [55] and the published literature, and supplemented with our own data and those of collaborators. For *T. elliptica*, we combined occurrence data for all species in the complex. A total of 852 observations were recorded for *T. elliptica* and 639 for *A. latifolia*. To avoid any potential biases that could arise as a result of geographic clustering of occurrence points, and, in turn, the over-representation of environmental conditions associated with areas of greater sampling effort, occurrence points were spatially thinned using the spThin package in R 4.3.2 [56] such that selected occurrence points were at least 10 km apart. After thinning, we had a total of 305 occurrence points for *T. elliptica* and 228 points for *A. latifolia*, with 56 grid cells (10×10 km) containing both species (Fig 1).

**Fig 1.**
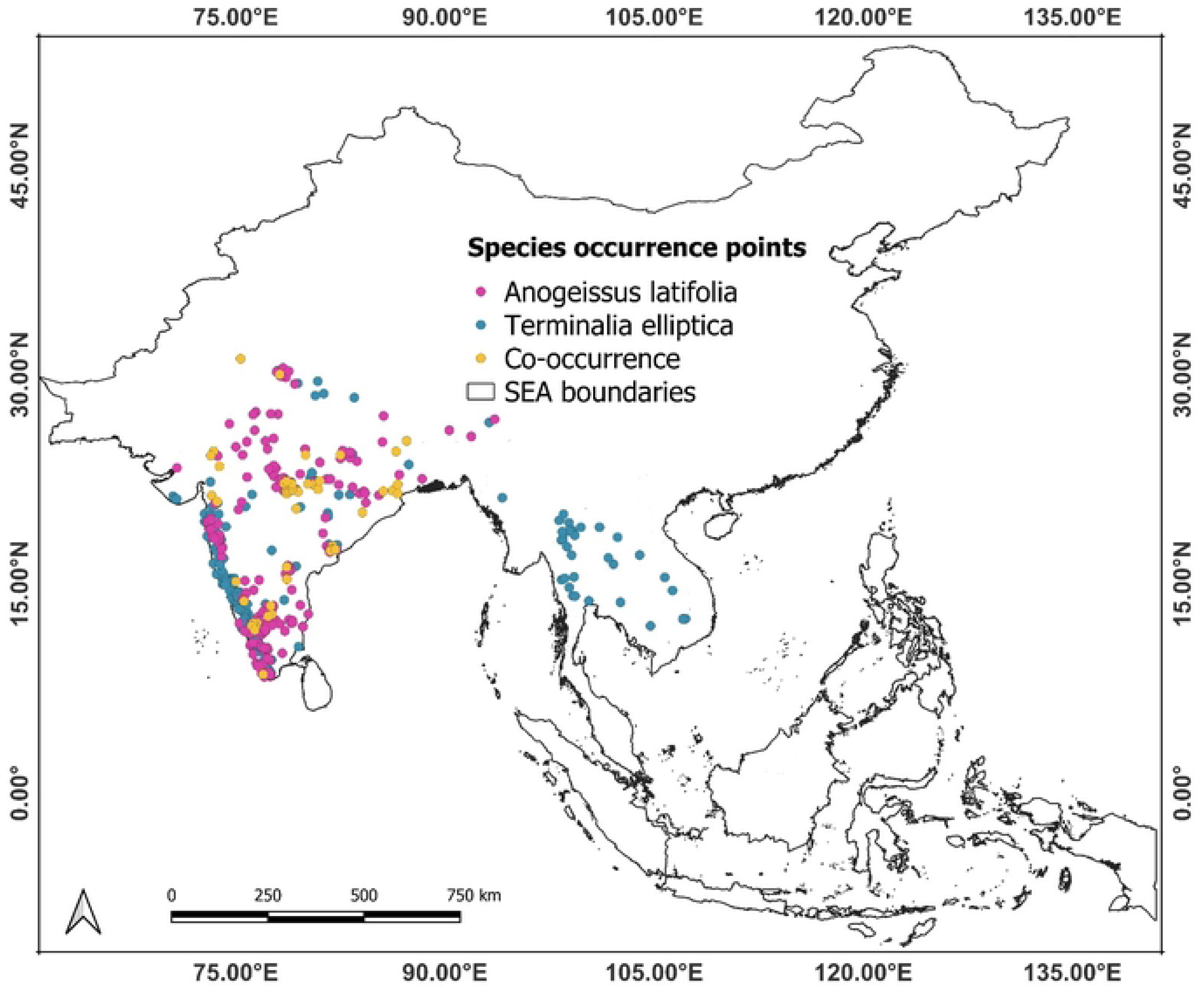
Map showing the location of species occurrence points used for the analyses. *A. latifolia* is represented in purple, *T. elliptica* in blue, and areas of species co-occurrence are marked in gold.

### 2.3 Climate variables

For the species distribution and niche quantification analysis, 19 climatic variables were obtained from the WorldClim version 2.1 database [57], representing a 30-year (1970-2000) average climate dataset at a resolution of 1 km (30 arc seconds). After testing for collinearity, we selected a subset of six ecologically relevant environmental variables, relating to temperature and moisture regimes that are known to exert strong controls on species distributions, for our analyses: annual mean temperature (AMT, Bio1), temperature annual range (TAR, Bio7), mean temperature of the driest quarter (MTDQ, Bio9), mean annual precipitation (MAP, Bio12), precipitation of the driest month (PDM, Bio14), and precipitation seasonality (PS, Bio15).

### 2.4 Niche quantification

Species niches were characterized using the Ecospat package in R [42], a tool that quantifies and compares the distinct and overlapping environmental ranges of two species. We quantified overlap in the environmental niche space of both species using Schoener’s *D* statistic, which ranges from 0 (no overlap in environmental space) to 1 (both species share the same environmental space; [58,59]). We assessed species niche equivalency [59] to determine whether the niches of both species were sufficiently similar to be considered equivalent and interchangeable (niche equivalency test). For this, species occurrences were pooled and randomly reallocated between the two species 1000 times while maintaining the number of occurrences for each species as in the original dataset. Based on these permutations, a null distribution of niche overlap (*D* values) was generated. If the observed value of niche overlap (*D*) was significantly lower than the overlap values from the null distribution, then the niches of the two species were considered non-equivalent (One-tailed test; [59]). We additionally used the niche similarity test to assess if the niches of the two species were more or less similar than expected by chance after accounting for available environmental conditions in the areas where the species are distributed [59,60]. In this case, we generated a null distribution of 1000 overlap values using randomly generated occurrences from across the geographic areas where both species are distributed. The observed value of niche overlap (*D*) was compared to the quantiles of the null distribution (two-tailed test) to determine if the ecological niches of the two species were more or less similar than expected by chance.

### 2.5 Species distribution modelling

Species distribution modelling (SDM) uses an associative or correlative approach to quantify the relationship between species and their environment. This approach combines species occurrence data with environmental variables to identify areas with similar climates and conditions that could potentially support the species with a certain probability [40]. We chose to model species distributions using Maxent [61], implemented in ENMeval 2.0 [62], which employs a presence-background method to predict species distributions [63,64].

For modelling, we used 303 occurrence points for *T. elliptica* and 226 for *A. latifolia*, along with 10,000 background locations selected probabilistically from a bias layer. The bias layer was derived from a model built using a pooled dataset of presence locations for both species and environmental variables from WorldClim [57] as predictor variables. Maxent habitat suitability predictions range from 0 to 1, with values below 0.5 indicating unsuitable habitat and values above 0.5 indicating potentially suitable habitat [61].

Separate models were built using ENMeval for each species using six different combinations of predictor transformations, known as feature classes (*LQH, LQ, QH, L, Q*, *H*- where *L* stands for linear, *Q* for quadratic, and *H* for hinge; [65,66]. Each combination was tuned using ten different regularization multiplier (RM) parameters (ranging from 0.5 to 5 in intervals of 0.5), which determines the penalty for including variables or their transformations in the model [65]. This tuning was performed to improve model fitting, as recommended for building simpler models with better transferability [67–69].

### 2.6 Model evaluation

We conducted a sequential model evaluation using a cross-validation approach to select the optimal combination of feature classes and regularization multipliers for the best-performing models. We prioritized models with the lowest average test omission rate and the highest average *AUC* (Area Under the Receiver Operator Curve). Test and training datasets were created using four geographically structured partitions (spatial blocks) [67]. The average evaluation metrics across partitions included measures of model transferability, such as *OR_MTP_* (omission rate of test presences in model predictions using a minimum training presence threshold) and *AUC_DIFF_* (difference between training and test Area Under the Receiver Operator Curve, which evaluates model overfitting). We also assessed model discriminatory ability using *AUC_TEST_*, which represents the probability that the model ranks a randomly selected presence location higher in habitat suitability than a randomly selected background location [69,70].

For each species, we compared models with *AUC_TEST_* > 0.6 sequentially, selecting those with the minimum OR_MTP_, followed by the minimum *AUC_DIFF_*, and finally the maximum *AUC_TEST_*[71] . The best-performing model for each species (S1 Table) was then used to generate predictions of habitat suitability across peninsular India. Spatial data were processed using the “rgeos” [72], “raster” [73], and “sp” [74] packages, while Maxent models were run using the “ENMeval” [75] and “dismo” [76] packages in R 4.3.2 [77].

### 2.7 Future projections

We used the best performing model for each species to predict future habitat suitability under the SSP3-7.0 (Shared Socioeconomic Pathways 3-7.0) scenario following the sixth assessment report of the Intergovernmental Panel on Climate Change (IPCC) for two periods: the short-term (2041-2060) and the long-term (2081-2100). SSP3-7.0 represents a scenario with medium to high emissions, where temperatures are projected to rise steadily by 2.1°C by 2041-2060 and by 3.5°C to 4°C above pre-industrial levels by 2100, with CO_2_ emissions expected to nearly double. This scenario assumes significant emissions due to limited climate policies, slow adoption of clean technologies, and continued high fossil fuel use. It also envisions fragmented societies, regional conflicts, uneven economic development, and ineffective global governance in addressing climate change. For future projections, three General Circulation Models (GCMs) from the CMIP6 framework were used: the Goddard Institute for Space Studies (GISS-E2-1-G), the Model for Interdisciplinary Research on Climate (MIROC6), and the Meteorological Research Institute Earth System Model (MRI-ESM2-0). These GCMs were selected based on their differing input layers and algorithms [78]. Future habitat suitability for both species was estimated based on the mean habitat suitability under the three GCMs. We compared modelled future projections with current modelled distributions to estimate shifts in the extent of suitable habitat for both species, identifying areas of potential gains and losses in the future.

## 3. RESULTS

### 3.1 Environmental niches of *T. elliptica* and *A. latifolia*

*T. elliptica* and *A. latifolia* show a high degree of niche overlap (Schoener’s D = 0.84), with the niche of *A. latifolia* appearing as a subset of *T. elliptica* (Figs 2A and B). Results of the one-tailed niche equivalency test indicate that the niches of the two species are equivalent (P = 1.0, Fig 2C), while the niche similarity test revealed that the niches of the two species are more similar than expected by chance (P< 0.05, Fig 2D). These results are also supported by the predicted niche occupancy profiles which indicate largely similar climatic tolerances for both species (Fig 3). Although *T. elliptica* occupied a wider rainfall range and occurred in wetter areas and those with cooler dry seasons (lower MTDQ) and lower temperature annual ranges than *A. latifolia* (Figs 3A and B), the occurrence densities of both species were largely overlapping for all the other environmental variables considered (Figs 3C-F).

**Fig 2.**
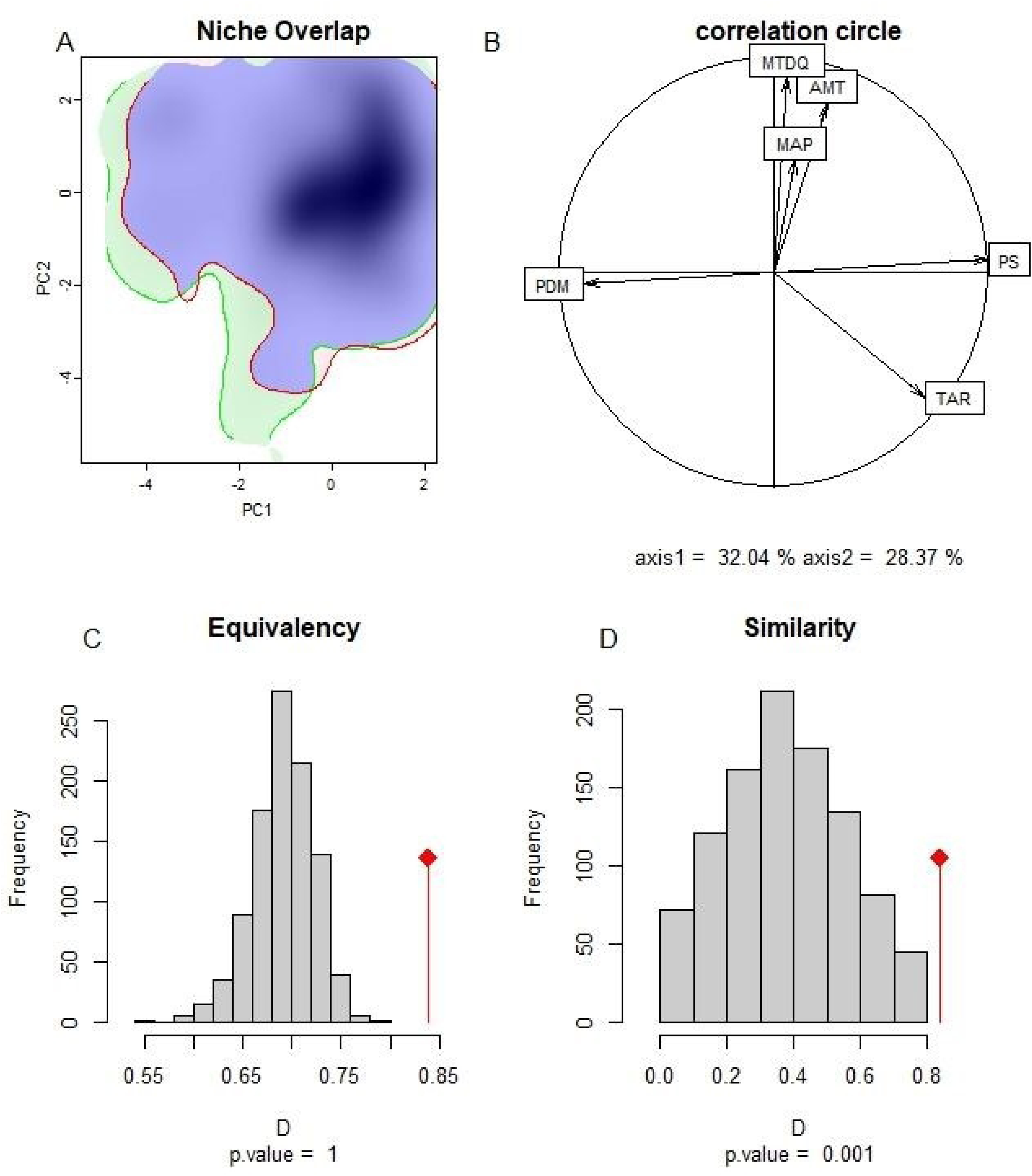
Climatic niche analyses of *T. elliptica* and *A. latifolia*. A) Visualization of niches of both species in PCA space, with the niche of *T. elliptica* shown in green, *A. latifolia* in red, and regions of niche overlap in blue. B) Correlation circle depicting the contribution of different environmental variables to the loading of the PCA axes. The first two principal components accounted for 60.41% of the variance in the selected environmental variables (PC1 = 32.04%, PC2 = 28.37%); MAP-mean annual precipitation, MTDQ-mean temperature driest quarter, TAR-temperature annual range, AMT-annual mean temperature, PS-precipitation seasonality, PDM-precipitation driest month. Panels C and D show the observed values of Schoener’s index of niche overlap (*D*) relative to the expected values of null distribution (*D*) for the tests of niche equivalency (C) and niche similarity (D).

**Fig 3.**
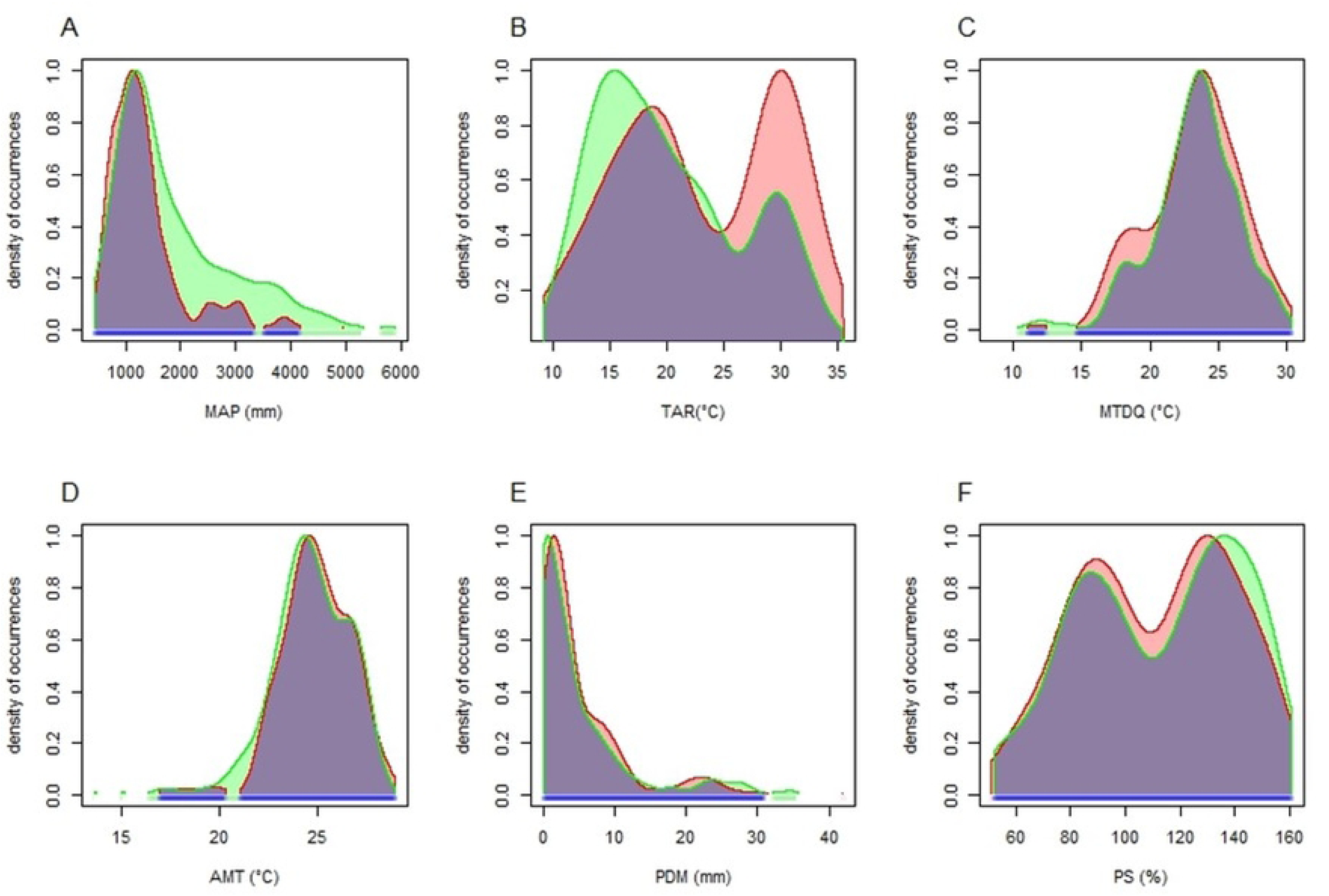
Predicted niche occupancy of *T. elliptica* and *A. latifolia* with respect to the different climatic variables included in the study (A) mean annual precipitation (MAP, Bio12), (B) temperature annual range (TAR, Bio7), (C) mean temperature of the driest quarter (MTDQ, Bio9), (D) annual mean temperature (AMT, Bio1), (E) precipitation of the driest month (PDM, Bio14), and (F) precipitation seasonality (PS, Bio15). Green and red shaded areas represent the occurrence densities of *T.elliptica* and *A.latifolia,* respectively, while blue areas indicate regions of niche overlap.

### 3.2 Current distributions of *T. elliptica* and *A. latifolia*

Habitat suitability maps, generated using Maxent, indicate a wide distribution of potentially suitable areas for both species across the subcontinent under current climatic conditions (Figs 4A and D). Areas of high suitability for both *A. latifolia* and *T. elliptica* include the Western Ghats, parts of Saurashtra in the northwest, the central Indian highlands including the Vindhya and Satpura ranges, and the western parts of the Indo-Gangetic plains (Figs 4A and D). Additionally, the northern Eastern Ghats and parts of Northeast India also appear to be suitable for *A. latifolia* (Fig 4A). The total area of suitable habitat (estimated by multiplying raster pixel suitability values by the area of grid cells) under current environmental conditions for *A. latifolia* is 4,16,057 km², while that of *T. elliptica* is 4,01,544 km² (Table 2).

**Fig 4.**
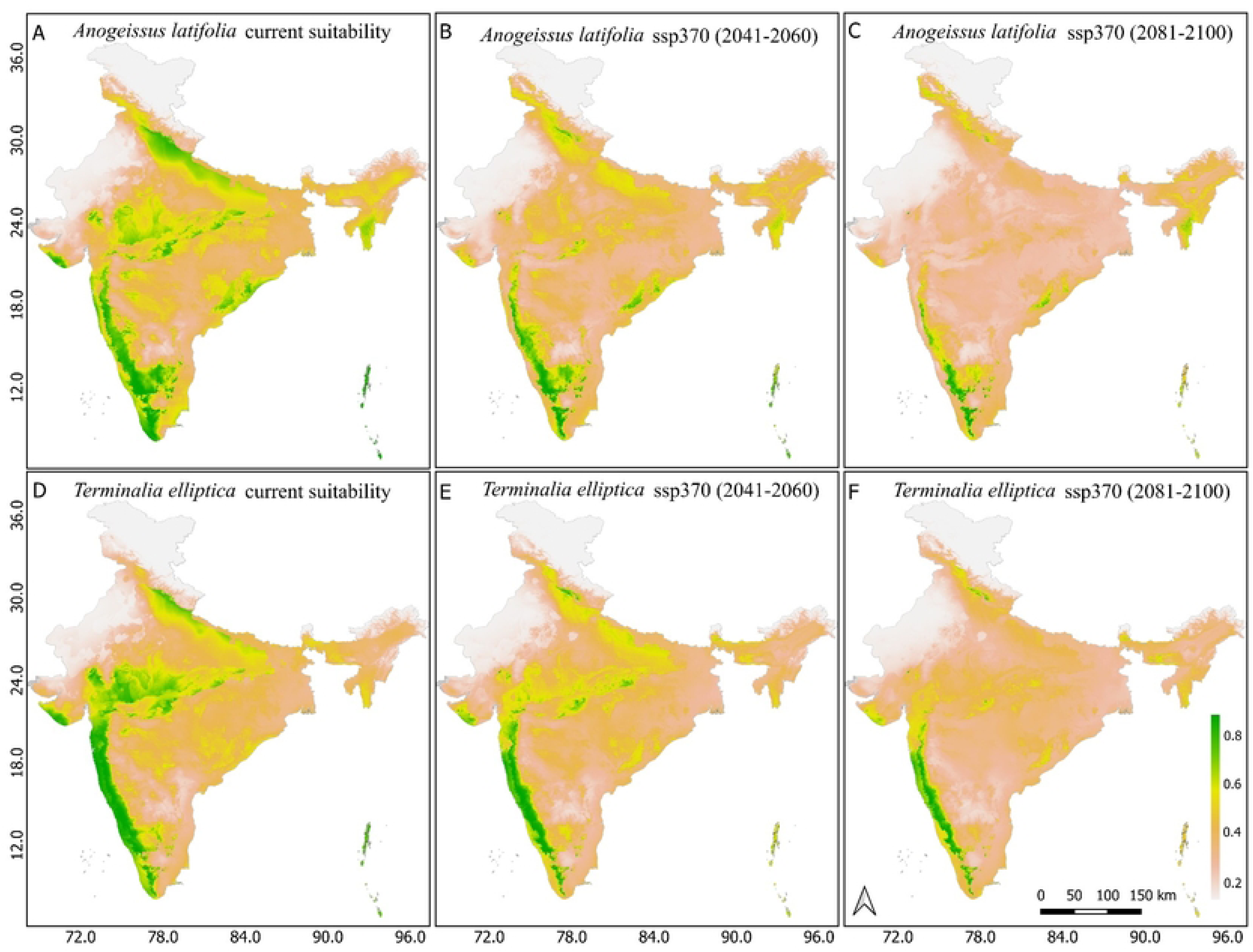
Current and future distribution of suitable habitat for *A. latifolia* and *T.elliptica*. Habitat suitability under current and future climates under the ssp370 scenarios for *A. latifolia* (A-C) and *T. elliptica* (D-F).

**Table 2.** Suitable habitat area (km^2^) for *T. elliptica* and *A. latifolia* under current and future (2041-60 and 2081-2100) climates under the ssp370 scenario (based on a habitat suitability cut-off of ≥ 70%).

| Species | Current<br>Suitable<br>habitat (km<sup>2</sup>) | Suitable habitat<br>(2041-60) (km<sup>2</sup>) | Suitable habitat<br>(2081-2100)<br>(km<sup>2</sup>) | Area loss<br>2041-2060<br>(km<sup>2</sup>)/ (%) | Area loss<br>2081-2100<br>(km<sup>2</sup>)/ (%) |
| --- | --- | --- | --- | --- | --- |
| *T. elliptica* | 4,01,544 | 1,69,334 | 1,08,054 | -2,32,210<br>(~57.8 %) | -2,93,490<br>(~73%) |
| *A. latifolia* | 4,16,075 | 1,52,773 | 6,5742 | -2,63,302<br>(~63%) | -3,50,333<br>(~84%) |

Despite having fairly similar niches, the predictor variables influencing the distribution of the two species differed (Fig 5). Mean annual precipitation (MAP), mean temperature of the driest quarter (MTDQ), temperature annual range (TAR) and annual mean temperature (AMT) were the primary variables influencing the distribution of *A. latifolia (*relative contributions of 42%, 24.18%, 21.65% and 11.76%, respectively), while precipitation seasonality (PS) and precipitation of the driest month (PDM) were largely unimportant (Fig 5). In contrast, MTDQ was the most important variable influencing the distribution of *T. elliptica* (relative contribution 38.3%), followed by TAR (18.3%), MAP (17.1%), PDM (8.92%), AMT (8.71%) and PS (8.65%) (Fig 5; S2 and S3 Tables).

**Fig 5.**
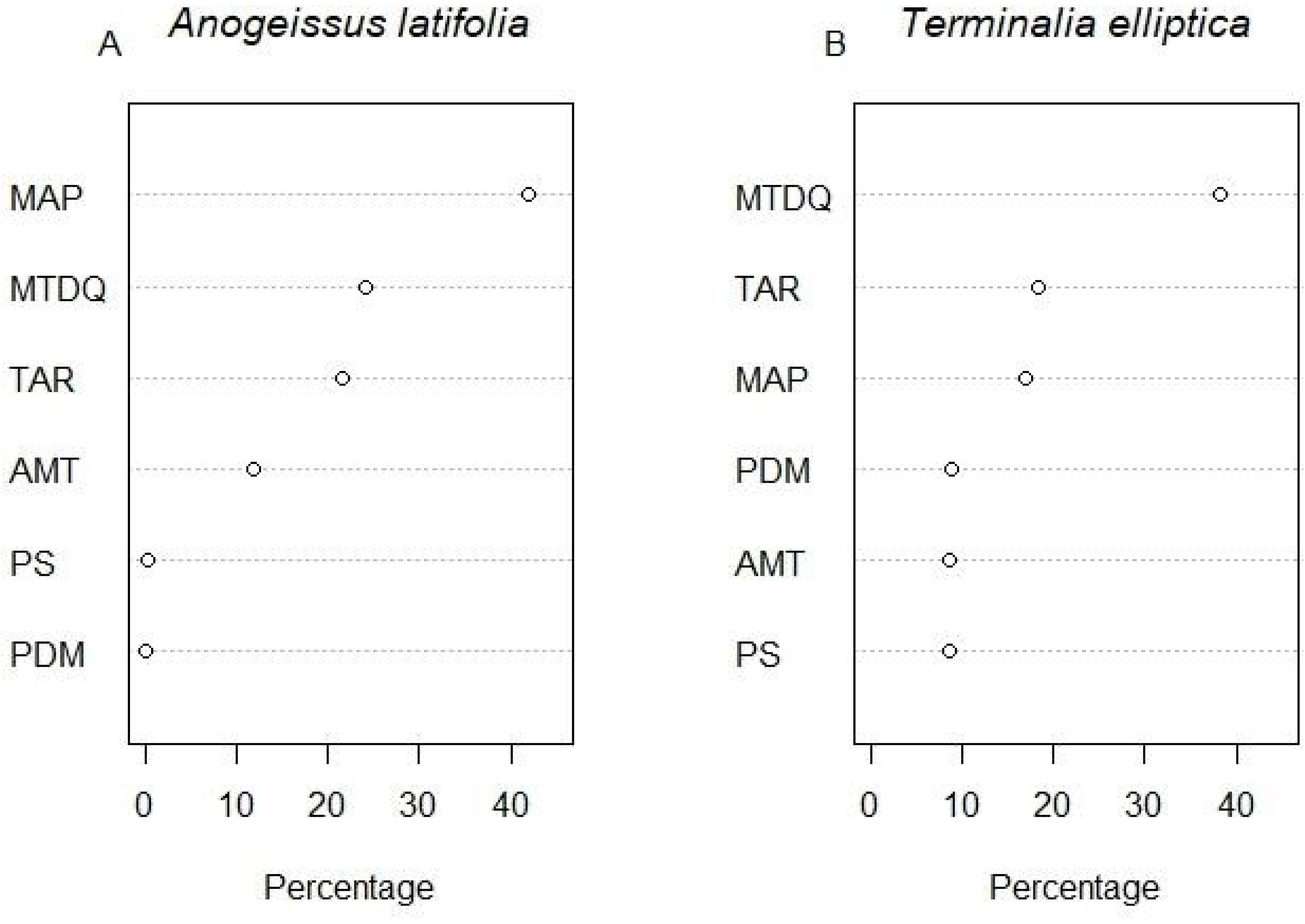
The contribution of different predictor variables to species habitat suitability of (A) *Anogeissus latifolia* and (B) *Terminalia elliptica*. AMT (Bio1) - annual mean temperature, TAR (Bio7) - temperature annual range, MTDQ (Bio9) - mean temperature of the driest quarter, MAP (Bio12) - mean annual precipitation, PDM (Bio14) - precipitation of the driest month, and PS (Bio15) - precipitation seasonality.

### 3.3 Species distributions under future climate conditions

Our models predict substantial reductions in the extent of suitable habitats for both *A. latifolia* and T*. elliptica* in the future under the SSP370 scenario (Figs 4 & 6). *A. latifolia* is predicted to lose ∼63 % of suitable habitat area by 2041-60 and ∼84% by the end of century (2081-2100; Figs 4 A-C). Similarly, *T. elliptica* is predicted to lose ∼57.8% and ∼73% of suitable habitat area by 2041-2060 and 2081-2100, respectively (Figs 4 D-F). Overall, both species are projected to lose large extents of currently suitable habitat, and only show modest gains in suitable habitats in the future - 4,288 km^2^ for *A.latifolia* and 3,072 km^2^ for *T.elliptica*, respectively, by 2100 (Fig 6 A-F, S2 Fig). For both species the most suitable habitats by the end of the century are likely to be concentrated in the Western Ghats, with most other parts of the country (e.g. central Indian highlands, Indo-Gangetic plains and the Eastern Ghats) witnessing substantial reductions in habitat quality for both species (Fig 4).

**Fig 6.**
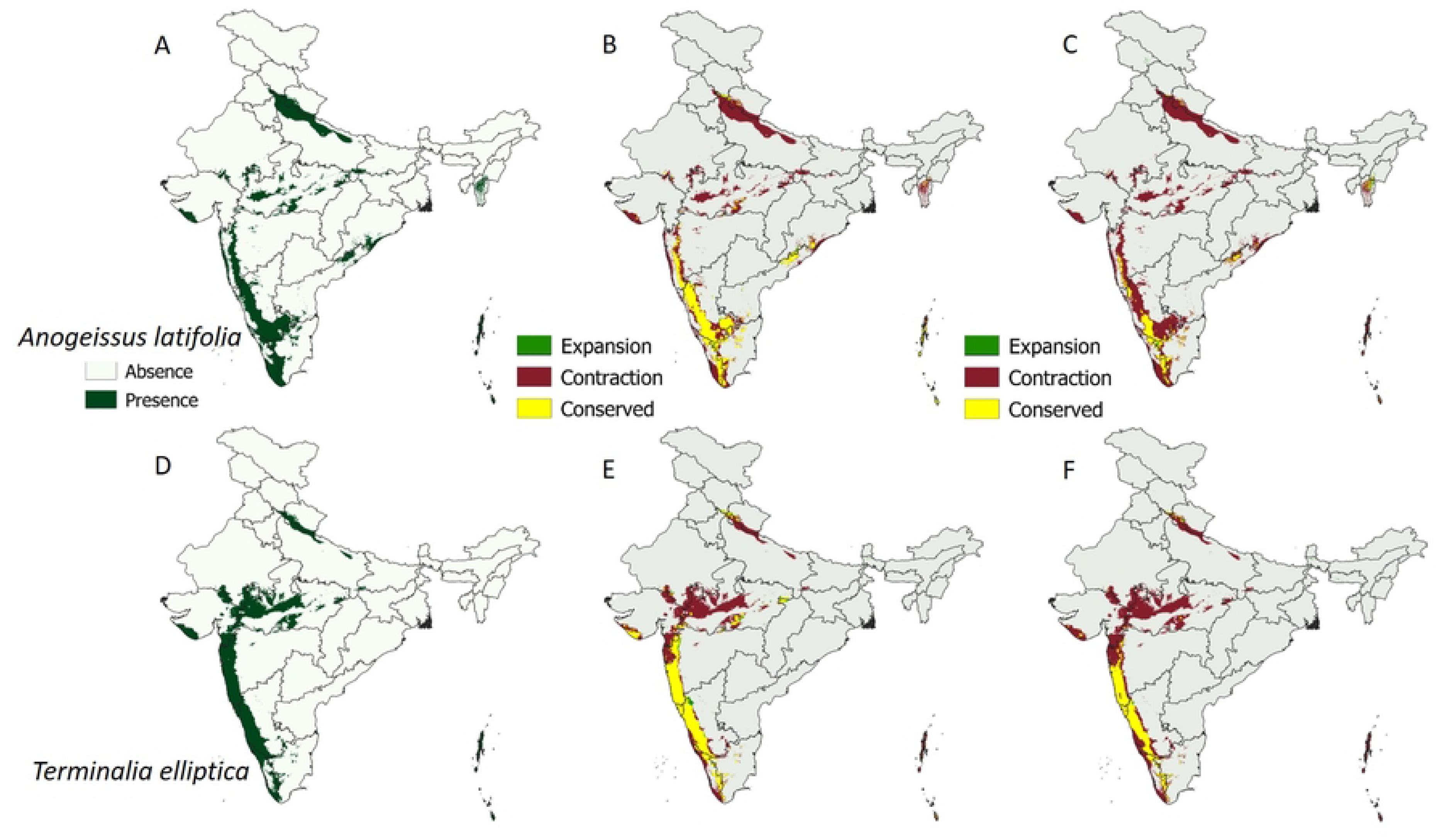
Extent of highly suitable habitat (≥ 70% suitability) for *A. latifolia* and *T. elliptica* under current conditions (A and D), and in the future under the SSP 370 scenario. Areas in yellow indicate regions where suitability remains unchanged, maroon indicates areas showing loss of suitability, while green indicates currently unsuitable regions that become suitable for *A. latifolia* (B and C) and *T. elliptica* (E and F) during the periods 2041–2060 and 2081– 2100, respectively.

The extent of overlap in the distributions of the two species when considering only locations with ≥ 0.7 probability of occurrence is predicted to decrease from 2,37,651 km2 currently to 84,444 km2 in 2041-60 to 39,283 km2 by 2081-2100 (Fig 6 A-F, S1 Fig). By 2100, the areas of co-occurrence of both species are projected to be largely restricted to the drier northern half of the Western Ghats (Fig 6 A-F, S1 Fig).

## 4. DISCUSSION

Our study species, *Anogeissus latifolia* and *Terminalia elliptica,* characteristic trees of the broad-leaved dry forests and mesic savannas of the Indian subcontinent, co-occur across a wide geography, from the arid northwest to the central Indian highlands in the east, across the Gangetic plains, and south into the peninsula and the Western Ghats. Under current climatic conditions, large areas of the subcontinent thus appear suitable for both species (*A. latifolia* - 4,16,057 km²; *T. elliptica-* 4,01,544 km²). However, both these currently common species appear to be particularly sensitive to anticipated future changes in climate and their ranges are projected to shrink dramatically by the turn of the next century (S1 Fig; Table 2).

Under future climate scenarios, *A. latifolia* is projected to lose up to four-fifths of its suitable range (∼63 % by 2041-60 & ∼84% & 2081-2100) while *T. elliptica* is projected lose up to three-fourths (∼57.8 % by 2041-60 & ∼73% & 2081-2100). Although the climate niches of both species are very similar (84% overlap, Fig 2), with mean annual precipitation (MAP), mean temperature of the driest quarter (MTDQ) and temperature annual range (TAR) being primary drivers, the relative importance of these drivers differs between species. Specifically,

*T. elliptica* appears to occupy a slightly wider climatic niche than *A. latifolia*, extending into wetter regions, areas characterized by cooler dry seasons (lower MTDQ) and lower annual temperature ranges (Figs 3A and B). These subtle differences appear to give *T.elliptica* a slight advantage, allowing it to retain a slightly greater proportion of suitable habitat under future scenarios where large tracts of dry regions in the subcontinent are predicted to become wetter [8,79]. Notably, future projections indicate that the Western Ghats may serve as key refugia for both species as habitat suitability across much of the remaining subcontinent is predicted to decline (Figs 4 and 6).

Given the similarities in their climatic niches, it is not surprising that both species exhibit largely congruent distributions at the country-wide scale with a fair amount of overlap (Figs 4 and S1 Fig), [80]. However, intriguingly, both species also co-occur at local scales and form co-dominant stands without one species displacing the other. Such co-existence hints at fine-scale resource partitioning along niche dimensions not considered in this study. However, a previous study reported very similar edaphic and topographic niches for both species even at neighbourhood scales, as well as very similar regeneration niches [80]. While these data suggest convergent adaptations of both species to commonly encountered environmental conditions, the exact mechanisms allowing them to coexist at local scales is not known. Further, whether such mechanisms also influence the current, and potential future distributions, of both species remains unknown.

Ultimately, the realized geographic distributions of the two species result not only from their climatic niches, but also other factors such as life history strategies, the capacity to tolerate and regrow after disturbances such as fire, drought, and herbivory, and the ability to regenerate naturally from seeds [20,32,35,50,81]. Worryingly, both species have been reported to be highly inconsistent and variable in their natural regeneration patterns from multiple sites across their range [34,35,50,80]. For example, although regeneration from seeds in *A.latifolia* appears to be consistent across years in some sub-Himalayan regions, it seems to be highly variable across years in some southern peninsular regions, with fertile seeds being produced only sporadically [37,50]. At a long-term monitoring site in southern India where the two species collectively accounted for ∼25% of all adult individuals in an area of fifty hectares, only 14 to 16 seedlings of each species recruited into the community over a 20-year period, most of which occurred during a single year [80]. The interactions of such life history traits with future climate regimes may further modify the expected range distributions for these species in the future.

The results of this study indicate that *A. latifolia* and *T. elliptica*, species that occur widely and form repeatable associations across large parts of the country, are likely to witness significant range reductions by the end of the century, with areas of future co-occurrence largely restricted to the northern portion of the Western Ghats (S1 Fig). What is presently unclear, however, is the extent to which the loss of these dominants will impact ecosystem functioning and the kind of vegetation formations that will replace the current *Anogeissus-Terminalia* associations. Future work aimed at identifying factors that currently limit regeneration of these two dominants across large parts of their range, and understanding how their loss is likely to impact subsequent community assembly will be critical for developing strategies to more effectively manage large tracts of broad-leaved deciduous forests and mesic savannas in the long-term.

## ACKNOWLEDGMENTS

We would like to acknowledge Abhirami Ravichandran, Ajith Ashokan, Amit Kurien, Aaroha Malagi, Dr. Ayesha Prasad, Dharmendra Khandal, Hansraj Gautam, Muhammed Maaz, Mukta Mande, Navendu Page, Rajat Rastogi, Siddarth Machado, Snehalatha, Vinay Sagar, the LEMoN-India network and Season watch for sharing species occurrence data. We also want to thank the National Centre for Biological Sciences (NCBS, TIFR) for providing access to their high-performance cluster to run the model. KL also acknowledges the fellowship support provided by CSIR-UGC and the R. M. Tulpule Charitable Trust.

